# Discovery of volatile biomarkers of Parkinson’s disease from sebum

**DOI:** 10.1101/469726

**Authors:** Drupad K Trivedi, Eleanor Sinclair, Yun Xu, Depanjan Sarkar, Camilla Liscio, Phine Banks, Joy Milne, Monty Silverdale, Tilo Kunath, Royston Goodacre, Perdita Barran

**Affiliations:** Manchester Institute of Biotechnology, School of Chemistry, The University of Manchester, Princess Street, Manchester, UK, M1 7DN; Anatune, 4 Wellbrook Way, Girton, Cambridge, UK, CB3 0NA; Department of Neurology, Salford Royal Foundation Trust, Manchester Academic Health Science Centre, University of Manchester, UK; Institute for Stem Cell Research, School of Biological Sciences, The University of Edinburgh, Edinburgh UK, EH16 4UU

## Abstract

Parkinson’s disease (PD) is a progressive, neurodegenerative disease that presents with significant motor symptoms, for which there is no diagnostic test (*1–3*). We have serendipitously identified a hyperosmic individual, a ‘Super Smeller’ that can detect PD by odor alone, and our early pilot studies have indicated that the odor was present in the sebum from the skin of PD subjects(*4*). Here, we have employed an unbiased approach to investigate the volatile metabolites of sebum samples obtained non-invasively from the upper back of 64 participants in total (21 controls and 43 PD subjects). Our results, validated by an independent cohort, identified a distinct volatiles-associated signature of PD, including altered levels of perillic aldehyde and eicosane, the smell of which was then described as being highly similar to the scent of PD by our ‘Super Smeller’.

**1 sentence summary:** Metabolomics identifies volatile odorous compounds from patient sebum that associate with the smell of Parkinson’s.

---

Physicians in ancient times, including Hippocrates, Galenus, and Avicenna, used odor as a diagnostic tool. Although the olfactory skills of physicians are not routinely used in modern medicine, it is well documented that a number of conditions, predominantly metabolic and infectious diseases, are associated with a unique odor(*5*), but there is scant evidence for odors as symptoms of neurodegenerative disorders. Joy Milne, a Super Smeller whose husband Les was diagnosed with PD in 1986, has demonstrated a unique ability to detect PD by odor(*4*). Joy has an extremely sensitive sense of smell, and this enables her to detect and discriminate odors not normally detected by those of an average olfactory ability. Preliminary tests with t-shirts and medical gauze indicated the odor was present in areas of high sebum production, namely the upper back and forehead, and not present in armpits(*4*). Sebum is a waxy, lipid-rich biofluid excreted by the sebaceous glands in the skin, over-production of which known as seborrhea, is a known non-motor symptom of PD(*6, 7*). Parkinson’s skin has recently been shown to contain phosphorylated **α**-synuclein, a molecular hallmark of PD (*8, 9*). Identification and quantification of the compounds that are associated with this distinctive PD odor could enable rapid, early screening of PD as well as provide insights into molecular changes that occur as the disease progresses and enable stratification of the disease in future.

Volatile organic compounds (VOCs) are often associated with characteristic odors, although some volatiles may also be odorless. The term ‘volatilome’ describes the entirety of the volatile organic and inorganic compounds that may originate from any organism, or object, which may be analytically characterized. For any given sample under ambient conditions in a confined environment, collecting, identifying, and measuring molecules in its headspace will then define its volatilome. Such measurements can be performed with thermal desorption gas chromatography mass spectrometry (TD-GC-MS), where a sample is placed in a closed vessel. The sample is then heated to encourage the production of volatiles and the headspace is captured for analysis by GC-MS. Investigation of volatile metabolites using mass spectrometry has proven to be extremely useful in clinical studies (*10–13*) as well as in the analysis of the consistency and provenance of edible items (*14–16*). Recently, TD-GC-MS has been used as a volatilome analysis platform for the detection of compounds from bacteria implicated in ventilator associated pneumonia(*12*), for differentiation between odors due to human and animal decomposition(*17*), as well as aerosol detection of the fumes from e-cigarettes(*18*). This versatility of TD-GC-MS for samples from many sources renders it highly suitable for use in identifying the metabolites that give rise to the distinct scent of PD. We have established a workflow that starts in clinics with the collections of sebum samples from the upper backs of PD patients along with matched control subjects and progresses to the discovery of disease specific volatile metabolites, the odor of which is confirmed by our Super Smeller. (Figure 1, Supplementary information and Table S1A).

**Figure 1:**
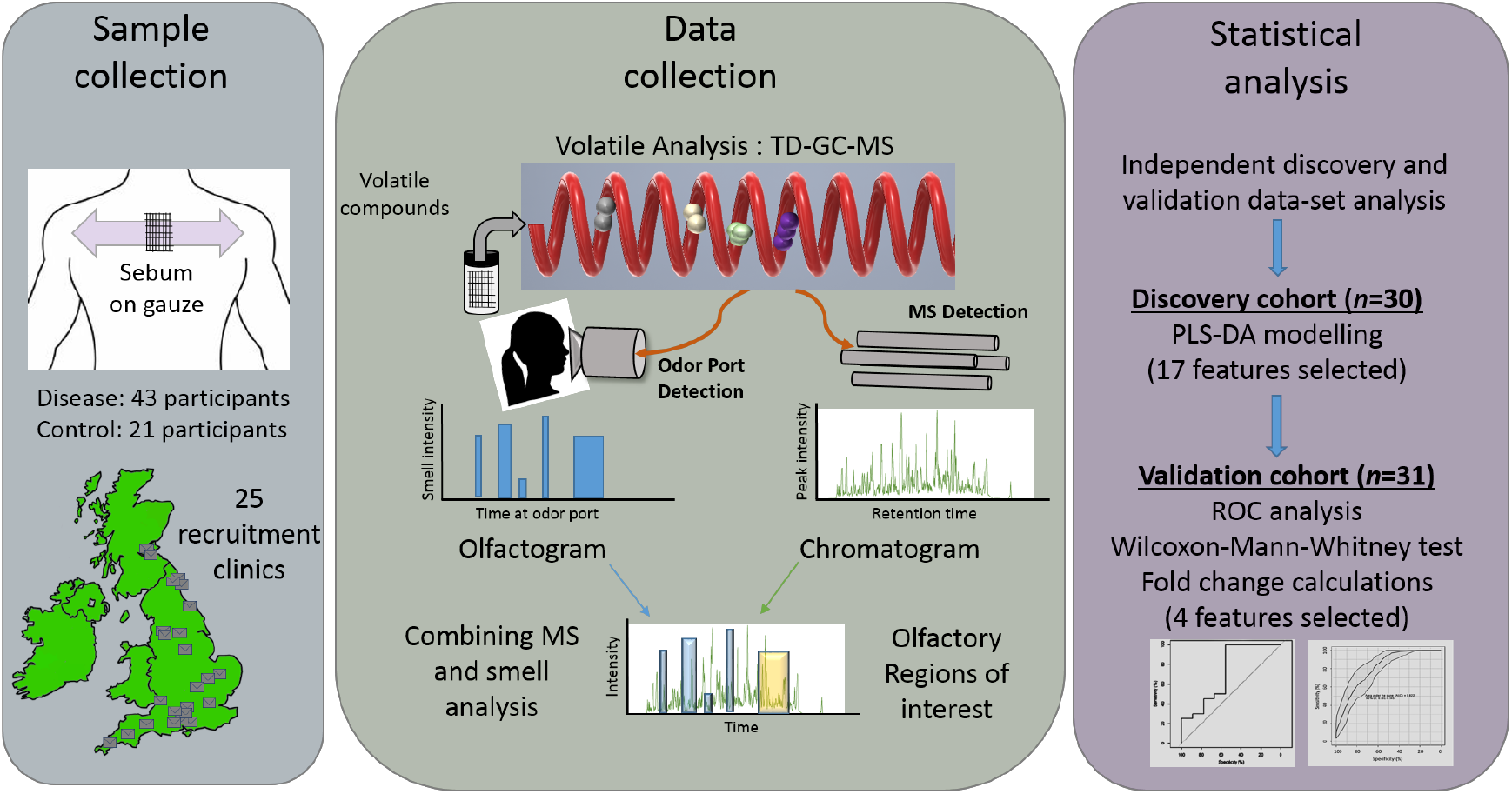
Schematic outline of the workflow described in this study - from sample collection to biomarker discovery. Parkinson’s disease patient samples and control participant samples were collected from 25 sites across the UK using gauze swabs to sample the sebum from the top back region from 64 people. Thermal desorption-gas chromatography-mass spectrometry (TD-GC-MS) analysis was performed alongside olfactory analysis results of which were then combined. Statistical analysis was performed on two independent cohorts. Data from discovery cohort consisting of 30 participants were used to create a partial least squares-discriminant analysis (PLS-DA) model and differential features found as a result were then targeted for presence in a separate validation cohort consisting of 31 participants. The significance of these biomarkers was tested using receiver operating characteristic (ROC) analyses and Wilcoxon-Mann-Whitney test. Finally, four features that showed similar statistical significance and expression on both cohorts were selected for biological interpretation of data.

In the current study, VOCs from the sample headspace were measured in two cohorts: a ‘discovery’ cohort and a ‘validation’ cohort, to validate discovered biomarkers (*19*) (for demographics see Table S1B). A third cohort consisting of three drug-naïve PD participants was used for mass spectrometry analysis in conjunction with a human Super Smeller *via* an odor port (Figure 1). This proof of principal study provides the first description of the skin volatilome in Parkinson’s disease compared to control subjects.

Mass spectrometry data were collected, deconvolved and pre-processed as described in the supplementary information. A partial least squares discriminant analysis (PLS-DA) model was built using the discovery cohort data (Figure 2). The classification accuracy of this model was validated by bootstrapping approach (n=1000). The variables, contributing to classification (n=17) were selected using variable importance in projections (VIP) scores where VIP > 1. We note at this stage, that one of the 17 metabolites found is 3,4-dihydroxy mandelic acid a metabolite of norephinephrine in humans. This catechol is also a metabolite of L-dopa, one of the most commonly prescribed medication for Parkinson’s, since here it is observed in drug naïve and control participants its presence may originate from endogenous mandelic acid. Norepinephrines including 3,4-dihydroxy mandelic acid are key molecules in the anabolism of brain neurotransmitters. Changes in neurons and neurotransmitters for instance, decrease of dopamines, a precursor to 3,4-dihydroxy mandelic acid is an extremely well-known characterization of PD(*20*).

**Figure 2:**
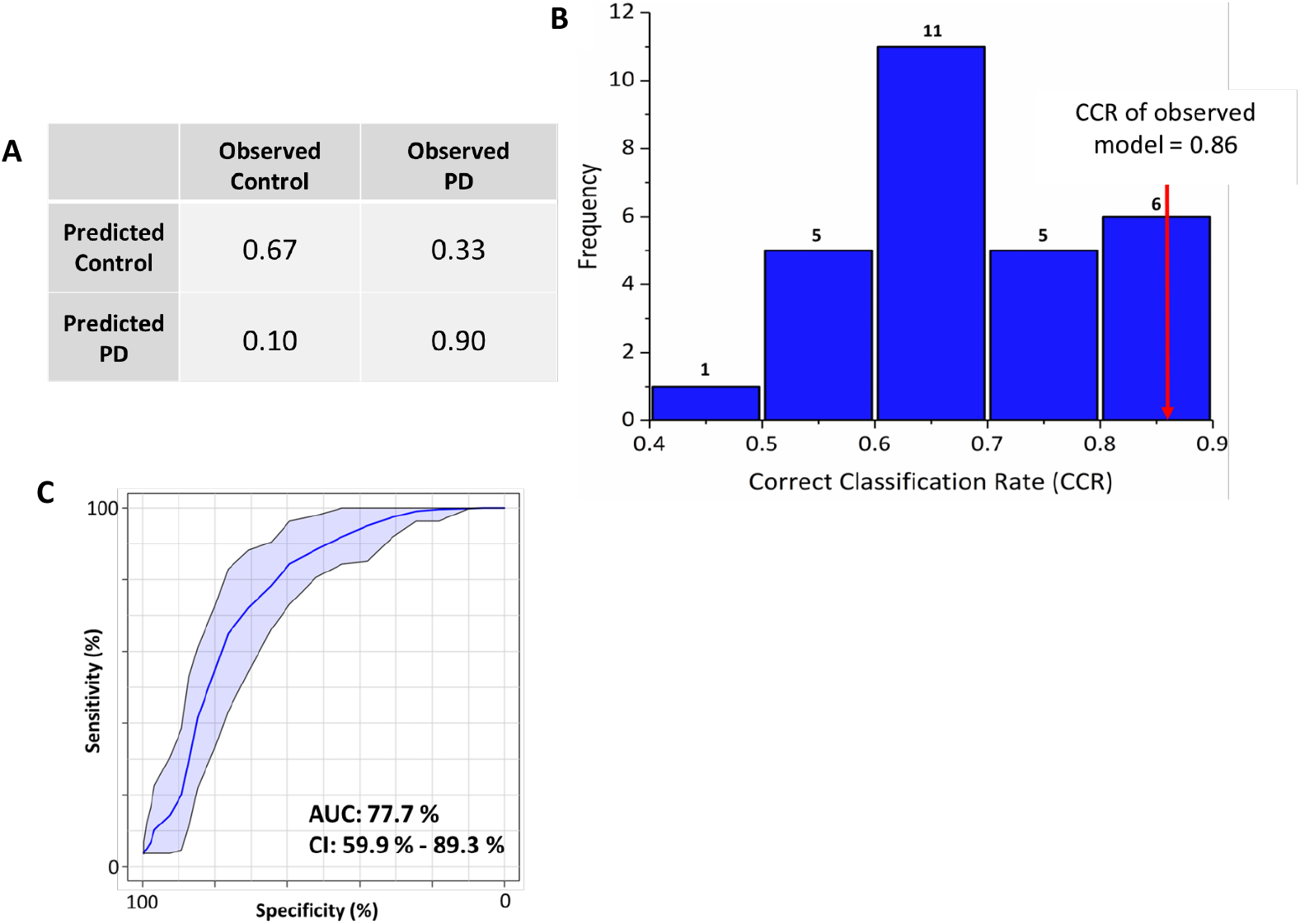
PLS-DA classification model. (A) Classification matrix of PLS-DA model validated using 5-fold cross validation showing 90% correct prediction of Parkinson’s disease samples (B) PLS-DA modelling was further tested using permutation tests (where the output classification was randomised; *n*=26) and results are plotted as a histogram which shows frequency distribution of correct classification rate (CCR) which yielded CCRs ranging between 0.4 to 0.9 for permutated models. The observed model was significantly better than most of the permuted models (*p* < 0.1); shown by the red arrow. (C) ROC plot generated using combined samples from both cohorts and the panel of four metabolites that were common and differential between control and PD. The shaded blue area indicates 95% confidence intervals calculated by Monte Carlo Cross Validation (MCCV) using balanced sub-sampling with multiple repeats.

The measured volatilome in the validation cohort data (from a different population than the discovery cohort) was targeted for the presence or absence of these discovered biomarkers. Out of these 17 metabolites, 13 were also found in the validation cohort data, and nine of these had retention times that allowed us to confidently assign them as identical (Table S2). These nine biomarkers found in both cohorts were selected for further analysis and statistical testing. To evaluate the performance of these biomarkers, we conducted receiver operating characteristic (ROC) analyses with data from both the discovery cohort and the validation cohort (Figure S1). ROC curves and Wilcoxon-Mann-Whitney tests as well as fold-change calculations on individual metabolites shows four out of these nine metabolites had a similar trend in regulation between the discovery and validation cohorts and their performance was also similar as measured by AUC (Table 1, Figure 3). We adhered to the Metabolomics Standards Initiative (MSI) guidelines for data analysis and for assignment of identity to features of interest(*19*) and all identified features are at MSI level two, which means these are putatively annotated compounds (e.g. without chemical reference standards, based upon physicochemical properties and/or spectral similarity with public/commercial spectral libraries). The compounds perillic aldehyde and eicosane are significantly different between PD and control in both the cohorts (*p* < 0.05): perillic aldehyde was observed to be lower in PD samples whereas eicosane was observed at significantly higher levels. Although hippuric acid and octadecanal were not significantly different (p > 0.05), the AUC and box plots (Figure 3) between the two cohorts were comparable and showed similar trends of being increased in PD.

**Figure 3:**
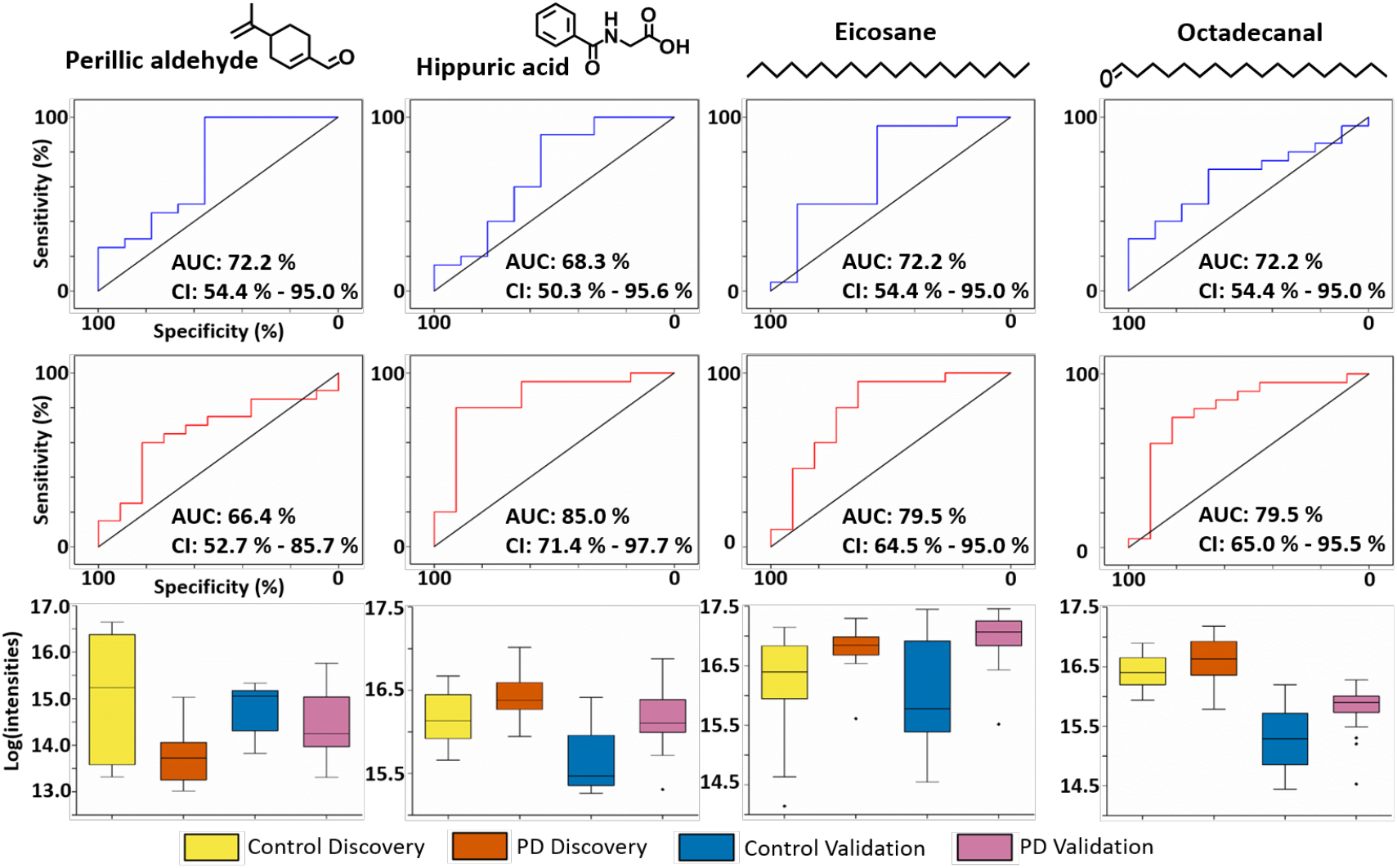
ROC curves and box plots for analytes of interest: In each panel from top to bottom: ROC curves for both discovery (blue) and validation (red) cohort for four analytes common to both experiments. Confidence intervals were computed with 2000 stratified bootstrap replicates and diagonal black line represents random guess. Box plots show comparison of means of log scaled peak intensities of these analytes, where black dots were outliers.

**Table 1:**
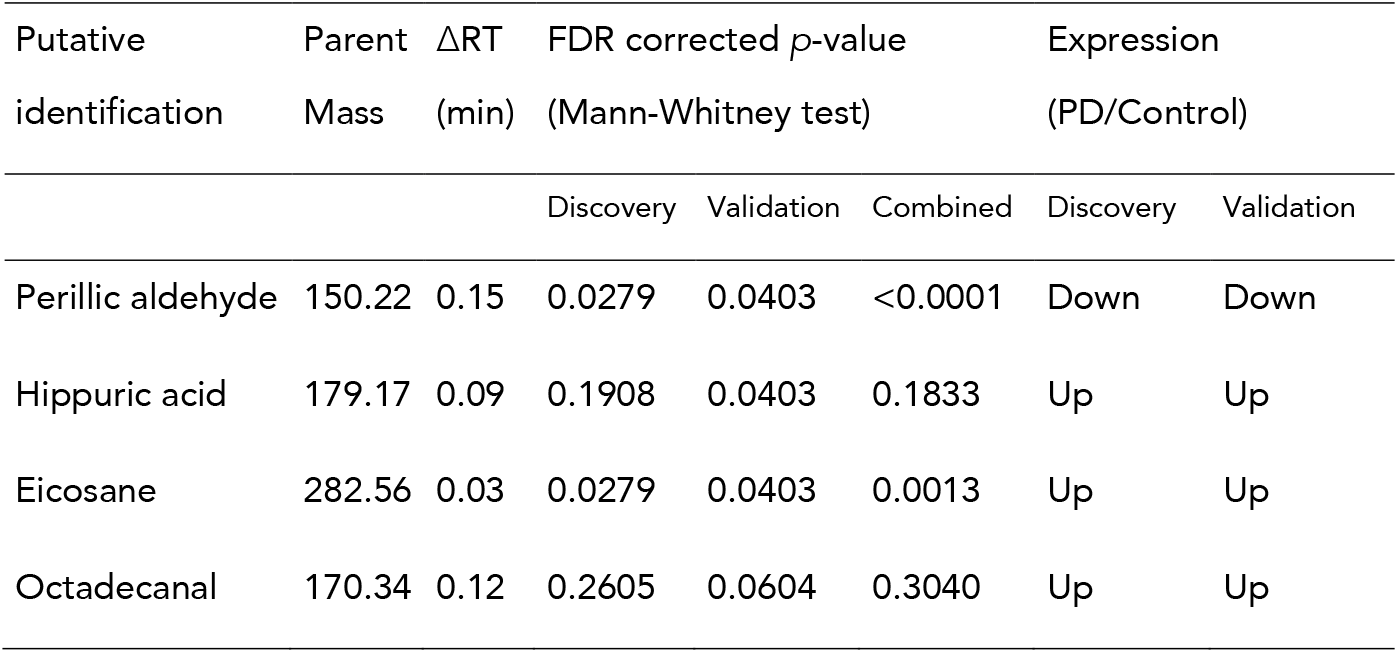
Panel of four volatile metabolites that were found to be differential between Parkinson’s and control samples, with similar trends observed in expression and AUC curves measured by ROC analyses. Perillic aldehyde and Eicosane were significantly down-regulated and up-regulated in PD, respectively (FDR corrected *p* < 0.05) in both cohorts.

| Putative<br>identification | Parent<br>Mass | ΔRT<br>(min) | FDR corrected $p$-value<br>(Mann-Whitney test) | | | Expression<br>(PD/Control) | |
| --- | --- | --- | --- | --- | --- | --- | --- |
|  |  |  | Discovery | Validation | Combined | Discovery | Validation |
| Perillic aldehyde | 150.22 | 0.15 | 0.0279 | 0.0403 | <0.0001 | Down | Down |
| Hippuric acid | 179.17 | 0.09 | 0.1908 | 0.0403 | 0.1833 | Up | Up |
| Eicosane | 282.56 | 0.03 | 0.0279 | 0.0403 | 0.0013 | Up | Up |
| Octadecanal | 170.34 | 0.12 | 0.2605 | 0.0604 | 0.3040 | Up | Up |

Using an odor port attached to the GC-MS instrument, the Super Smeller identified times at which any smell was present and also more importantly the times at which a specific ‘musky’ smell of PD was detected. Data were presented in the form of an olfactogram, where the presence and relative intensity of each smell were recorded at its corresponding chromatographic retention time. Olfactogram results obtained from the odor port were overlaid on the respective total ion chromatogram from GC-MS (Figure 4A). There was significant overlap between regions that contained up-regulated compounds and regions in which a smell similar or identical to that of PD scent was present. In the chromatographic trace the region between 19 and 21 min is of particular interest (Figure 4B) since the smell associated with the mixture of analytes in that window was described as “very strong” and “musky”. This is the same region where three compounds *viz*. hippuric acid, eicosane and octadecanal have been detected in both cohorts and all three were found to be up-regulated in PD subjects.

**Figure 4:**
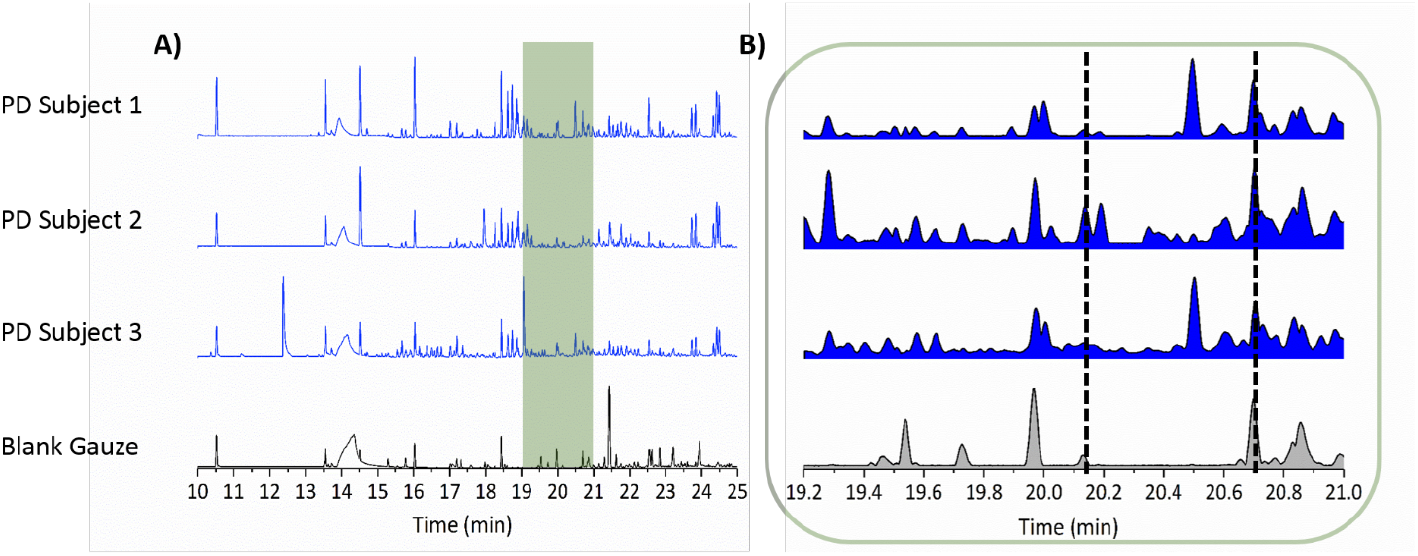
Comparison of GC-MS chromatogram to description of olfactory data described by the Super Smeller: GC-MS chromatogram from three drug naïve Parkinson’s subjects and a blank gauze. (A) The 10 to 25 min retention time range of the chromatographic analysis in which the Super Smeller described various odors associated with different GC-MS peaks. The overlaid green shaded area shows the overlap between real time GC-MS analysis and the Super Smeller describing a ‘strong PD smell’ via the odor port. (B) A zoom of the green highlighted area from A. This region is of particular interest as 3 out of 4 identified compounds are found here (Tables 1 and S2); it encompasses the time during which the Super Smeller described a musky PD-like scent as being ‘very strong’ (between the time lines at 19 and 21 min) for the PD samples and not for the blank. It can be noted that none of these compounds are found in blank gauze (bottom chromatogram) within the same retention time window as shown by normalized relative peak intensities to the highest peak in each chromatogram.

In order to validate mass spectrometry led biomarkers and to verify the resultant scent, the candidate compounds listed in Table S2 (n=17) were purchased and spiked onto gauze swabs (Table S3). An exploratory study with our Super Smeller was performed in which multiple mixtures of compounds (*n=5*) were spiked onto both blank gauze swabs and swabs that contained control sebum. Two final dispensed volumes of the mixtures were tested (40 μL and 100 μL) and all compounds used were at a single concentration (10 μM). In these blinded tests the Super Smeller grouped the samples in order of PD-like odor. She was able to isolate the swabs with a sebum background matrix and described them as more familiar to the PD-like smell than without control sebum. Further tests utilized control sebum as a background matrix for spiking candidate compounds and a range of concentrations was then selected for testing. Mixtures of the candidate compounds (*n*=17) were prepared at a range of concentrations (10 μM, 5 μM, 0.5 μM, 0.05 μM, 0.005 μM) and presented to the Super Smeller in a second blinded test, she was again asked to rank in order of PD-like smell. These results demonstrated she could detect (although not systematically order) the whole range of concentrations offered, and a concentration between 0.05 μM and 0.5 μM gave her the best response. A validation study consisting of three compound mixtures with significance from the MS analysis aimed to distinguish the combination that best gave rise to the most PD-like smell. Three mixture combinations were chosen at a single concentration (0.5 μM); all candidate compounds (*n=17*), all compounds identified in both the discovery and validation cohorts (*n=9*) and the panel of compounds expressed in same direction and differential between PD and control (*n=4*). The mixture of 9 compounds was consistently described as being most akin to the PD-like odor and was slightly overlapped by description and rank with the mixture of 4 compounds. The mixture of 17 compounds was grouped as the same ‘smell’ as the other two combinations however were described as significantly weaker. We hypothesize this is due to a lower concentration of each compound in the mixture and thus higher interference from background solvent smell. The results from these tests are depicted in Figure S2 whereby the intensity and correlation to the PD-like smell partitions the groups of samples tested. We do not conclude that these chemicals alone constitute the unique smell associated with PD, rather we demonstrate that they contribute to it.

From results obtained from three independent sets of data, from different people with one underlying factor (i.e. PD) separating them, it was clear that several volatile features were found to be significantly different between control and PD participants. There were no significant differences observed between PD participants on medication and drug naïve PD participants (p > 0.05 for all measured volatiles), indicating that the majority of the analysed volatilome and by inference sebum, are unlikely to contain drug metabolites associated with PD medication. In addition, applying machine learning approaches did not lead to a classification between drug naïve PD participants and PD participants on medication (Table S4). Perillic aldehyde and octadecanal are ordinarily observed as plant metabolites or food additives. It can be hypothesised that with increased and altered sebum secretion such lipid-like hydrophobic metabolites may be better captured or retained on the sebum-rich skin of PD subjects. Skin disorders in Parkinson’s have been observed previously and seborreheic dermatitis (SD) in particular has been flagged as a premotor feature of PD (*21*). It has been reported by Arsenijevic and coworkers (*7*) that PD patients who suffer from SD have increased *Malassezia* density on their skin and commensurate higher lipase activity required metabolically by yeast. This increased lipase activity could correlate with the enhanced production of eicosane, perillic aldehyde and octadecanal as highly lipophilic molecules since *Malassezia* requires specific exogenous lipids for growth. Eicosane is reported as being produced by *Streptomyces* as an anti-fungal agent (*22*) which also supports its increased presence on the skin of PD sufferers. The effects observed in our study could also signal altered microbial activity on the skin of PD patients that may affect the skin microflora causing changes in the production of metabolites such as hippuric acid (*23*). These potential explanations for the change in odor in PD patients all suggest a change in skin microflora and skin physiology that is highly specific to PD.

In conclusion, our study highlights the potential of comprehensive analysis of sebum from PD patients and raises the possibility that individuals can be screened non-invasively based on targeted analysis for these volatile biomarkers. We do acknowledge that the current study is limited with smaller sample size but the power of this study is a *different* validation cohort that consisted of completely different participants. This validation cohort was able to verify the findings and classification model built using data from our discovery cohort. A larger study with extended olfactory data from human smellers as well as canine smellers in addition to headspace analyses is the next step in further characterizing the PD sebum volatilome. This will enable the establishment of a panel of volatile biomarkers associated with PD and will open new avenues for stratification as well as facilitate earlier detection of PD and further the understanding of disease mechanisms.

## Supplementary information

includes description of sampling methodology, analytical methods used and has supplementary figures and tables as well as list of all the PIs from recruitment centres that participated in this study.

## Acknowledgment

We thank Michael J Fox Foundation and Parkinson’s UK for funding this study. We also thank our nationwide NIHR recruitment centers for their enthusiasm and rigor during the recruitment process. We are very grateful to all the participants who took part in this study as well as PIs and nurses across all the recruiting centers that helped us with the recruitment of participants.

## Author Contributions

DKT, ES, CL, JM and PB performed or contributed to the sample preparation, data collection. DKT and YX performed statistical analysis and validation of data. DKT, ES, JM, RG and PB designed the experiments. PB, RG, MS and TK conceived and supervised the study. DKT wrote the manuscript and all authors commented and contributed to the final version of manuscript.

## Materials and Correspondence

All materials and correspondence request should be addressed to Professor Perdita Barran at

